# Structure of nonstructural protein 1 from SARS-CoV-2

**DOI:** 10.1101/2020.11.03.366757

**Authors:** Lauren K. Clark, Todd J. Green, Chad M. Petit

**Affiliations:** Department of Biochemistry and Molecular Genetics, University of Alabama at Birmingham School of Medicine, Birmingham, Alabama 35294, USA; Department of Microbiology, University of Alabama at Birmingham School of Medicine, Birmingham, Alabama 35294, USA

**Keywords:** SARS-CoV-2, nonstructural protein 1, coronavirus, Severe Acute Respiratory Syndrome, X-ray crystallography

## Abstract

The periodic emergence of novel coronaviruses (CoVs) represents an ongoing public health concern with significant health and financial burden worldwide. The most recent occurrence originated in the city of Wuhan, China where a novel coronavirus (SARS-CoV-2) emerged causing severe respiratory illness and pneumonia. The continual emergence of novel coronaviruses underscores the importance of developing effective vaccines as well as novel therapeutic options that target either viral functions or host factors recruited to support coronavirus replication. The CoV nonstructural protein 1 (nsp1) has been shown to promote cellular mRNA degradation, block host cell translation, and inhibit the innate immune response to virus infection. Interestingly, deletion of the nsp1-coding region in infectious clones prevented the virus from productively infecting cultured cells. Because of nsp1’s importance in the CoV lifecycle, it has been highlighted as a viable target for both antiviral therapy and vaccine development. However, the fundamental molecular and structural mechanisms that underlie nsp1 function remain poorly understood, despite its critical role in the viral lifecycle. Here we report the high-resolution crystal structure of the amino, globular portion of SARS-CoV-2 nsp1 (residues 10 – 127) at 1.77Å resolution. A comparison of our structure with the SARS-CoV-1 nsp1 structure reveals how mutations alter the conformation of flexible loops, inducing the formation of novel secondary structural elements and new surface features. Paired with the recently published structure of the carboxyl end of nsp1 (residues 148 – 180), our results provide the groundwork for future studies focusing on SARS-CoV-2 nsp1 structure and function during the viral lifecycle.

**IMPORTANCE:** The Severe Acute Respiratory Syndrome Coronavirus 2 (SARS-CoV-2) is the causative agent for the COVID-19 pandemic. One protein known to play a critical role in the coronavirus lifecycle is nonstructural protein1 (nsp1). As such, it has been highlighted in numerous studies as a target for both the development of antivirals and for the design of live-attenuated vaccines. Here we report the high-resolution crystal structure of nsp1 derived from SARS-CoV-2 at 1.77Å resolution. This structure will facilitate future studies focusing on understanding the relationship between structure and function for nsp1. In turn, understanding these structure-function relationships will allow nsp1 to be fully exploited as a target for both antiviral development and vaccine design.

## INTRODUCTION

The periodic emergence of novel coronaviruses (CoVs) represents an ongoing public health concern with significant health and financial burden worldwide. In the past two decades, three highly pathogenic human CoVs have emerged from zoonotic events. In 2002, an outbreak of atypical pneumonia termed Severe Acute Respiratory Syndrome (SARS) appeared in the Guangdong Province of Southern China. The etiological agent for this disease was found to be a novel coronavirus named SARS coronavirus (SARS-CoV) (1–3). In 2012, a novel coronavirus named Middle East Respiratory Syndrome Coronavirus (MERS-CoV) was isolated from a patient who died in Saudi Arabia after presenting with acute respiratory distress and kidney injury.(4) The most recent occurrence originated in the city of Wuhan, China where a novel coronavirus, Severe Acute Respiratory Syndrome Coronavirus 2 (SARS-CoV-2), emerged causing severe respiratory illness and pneumonia (5, 6). The resulting disease was named Coronavirus Disease 2019 (COVID-19) and a pandemic was declared by the World Health Organization in March of 2020.

CoVs are members of the *Coronaviridae* family of viruses that contain large positive-sensed, single-stranded RNA genomes of approximately 30 kb in length. Approximately two-thirds of the genome, located in the 5’-genomic region, contains open-reading frame 1a (ORF1a) and open-reading frame 1b (ORF1b). To express both polyproteins, coronaviruses encode a slippery sequence and an RNA pseudoknot that cause a −1 ribosomal frameshift (7, 8). ORF1a and ORF1ab encode nsps 1 – 11 and 1 – 16, respectively, and are translated into large polyproteins which are subsequently cleaved into individual nonstructural proteins (nsps) by virally encoded proteases (9). Genes that encode the structural proteins (spike, envelope, membrane, and nucleocapsid) are located in the remaining third of the genome with a variable number of strain-dependent ORF accessory proteins present between the structural genes (10).

Nsp1 is the N-terminal cleavage product released from the replicase polyprotein by the virally encoded papain-like proteinase (nsp3d; PLpro) (11). It has been shown to promote cellular mRNA degradation, block host cell translation, and inhibit the innate immune response to virus infection (12–20). Deletion of the nsp1-coding region in infectious clones prevented the virus from productively infecting cultured cells (21). Also, mutations preventing the release of nsp1 from the nascent ORF1a polyprotein substantially limited virus viability (22). While there have been multiple studies on describing nsp1 function, there have been limited structure function studies focusing on nsp1. These types of studies are needed to fully exploit nsp1 as a target for both the development of antivirals and rational vaccine. The availability of high-resolution structures is a critical first step in establishing structure-function relationships of proteins and is the focus of this manuscript.

Here we report the high-resolution crystal structure of the globular portion of SARS-CoV-2 nsp1 to a resolution of 1.77 Å. A comparison of our structure with the SARS-CoV-1 nsp1 structure reveals how mutations alter the conformation of flexible loops. These changes in conformation induce the folding of novel secondary structural elements that, in turn, generate surfaces that are distinct from SARS-CoV-1 nsp1 in both contour and feature. Comparison with α-CoV nsp1 structures shows that the nsp1globular domain has a shared structural homology, while having low sequence similarity. Our results provide the groundwork for future studies focusing on the structure-function mechanisms by which SARS-CoV-2 nsp1 functions during the viral lifecycle.

## RESULTS

### Expression of SARS-CoV-2 nsp1

To obtain the high-resolution structure of nsp1 from SARS-CoV-2, we based our expression construct on the previously solved structure of nsp1 from SARS-CoV-1 (23). In this study, it was determined that nsp1 of SARS-CoV-1 includes a globular domain of residues 13 to 121 that is in between disordered regions consisting of resides 1 to 12 and 122 to 179. We generated an initial expression construct with N-terminal truncations beginning at amino acid residue 13 and continuing to residue 127. Using nuclear magnetic resonance spectroscopy, heteronuclear single quantum coherence triple resonance spectra analysis indicated that the amino terminus of our construct beginning at amino acid residue 13 prematurely eliminated a β-strand (data to be reported elsewhere). We generated a second expression construct with N-terminal truncations beginning at amino acid residue 10 continuing to residue 127. Both nsp1 constructs were expressed and purified to homogeneity.

### Crystal Structure of nsp1_10-127_ from SARS-CoV-2

Crystallization trials were initiated with nsp1_10-127_ and nsp1_13-127_. Crystals were obtained with each construct though the number of crystallization conditions were greater and crystal quality better with the longer construct, nsp1_10-127_. For this reason, structural studies were carried out with this construct. The structure of the globular domain of nsp1 was determined by molecular replacement with the core of SARS-CoV-1 nsp1 (PDB ID: 2HSX). Final data collection and refinement statistics are shown in Table 1. The final model contained residues 10-126 **(Fig. 1)**. The tertiary fold of nsp1 is composed of regular secondary structural elements arranged sequentially as β1–η1–α1–β2–η3–β3–β4–β5–β6–β7 with η1 and η2 being 3_10_ helices. A seven-stranded β–barrel is formed with a mixture of parallel and antiparallel β strands with contacts between β2, β3, β4, β5, and β6 strands as well as β1 and β7 strands. The β–strands consist of residues 13-20 (β1), 51-54 (β2), 68-73 (β3), 84-92 (β4), 95-97 (β5), 103-109 (β6), and 117-123 (β7). The η1 (residues 23-25) and η2 (residues 61-63) helices are positioned across one barrel opening while α1 (residues 34-49) is positioned alongside the barrel making contact with the β4 strand.

**FIG 1.**
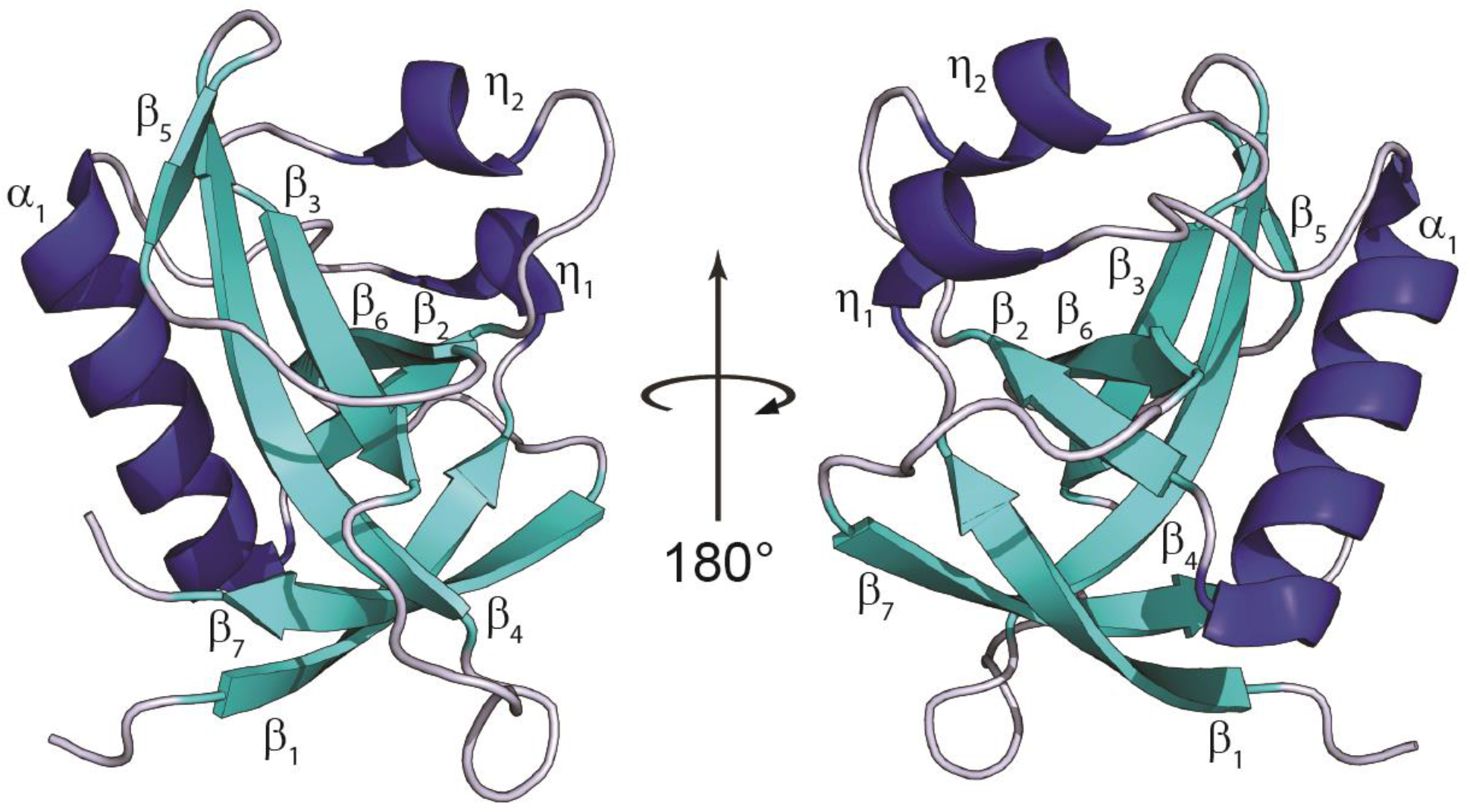
Crystal structure of SARS-CoV-2 nsp1. Ribbon diagram of the SARS-CoV-2 nsp1_10-126_ with the α-helices in blue, β-strands in cyan, and loops in grey. All secondary structural elements are labeled. Coordinates have been deposited under PDB ID: 7K7P

**Table 1.**
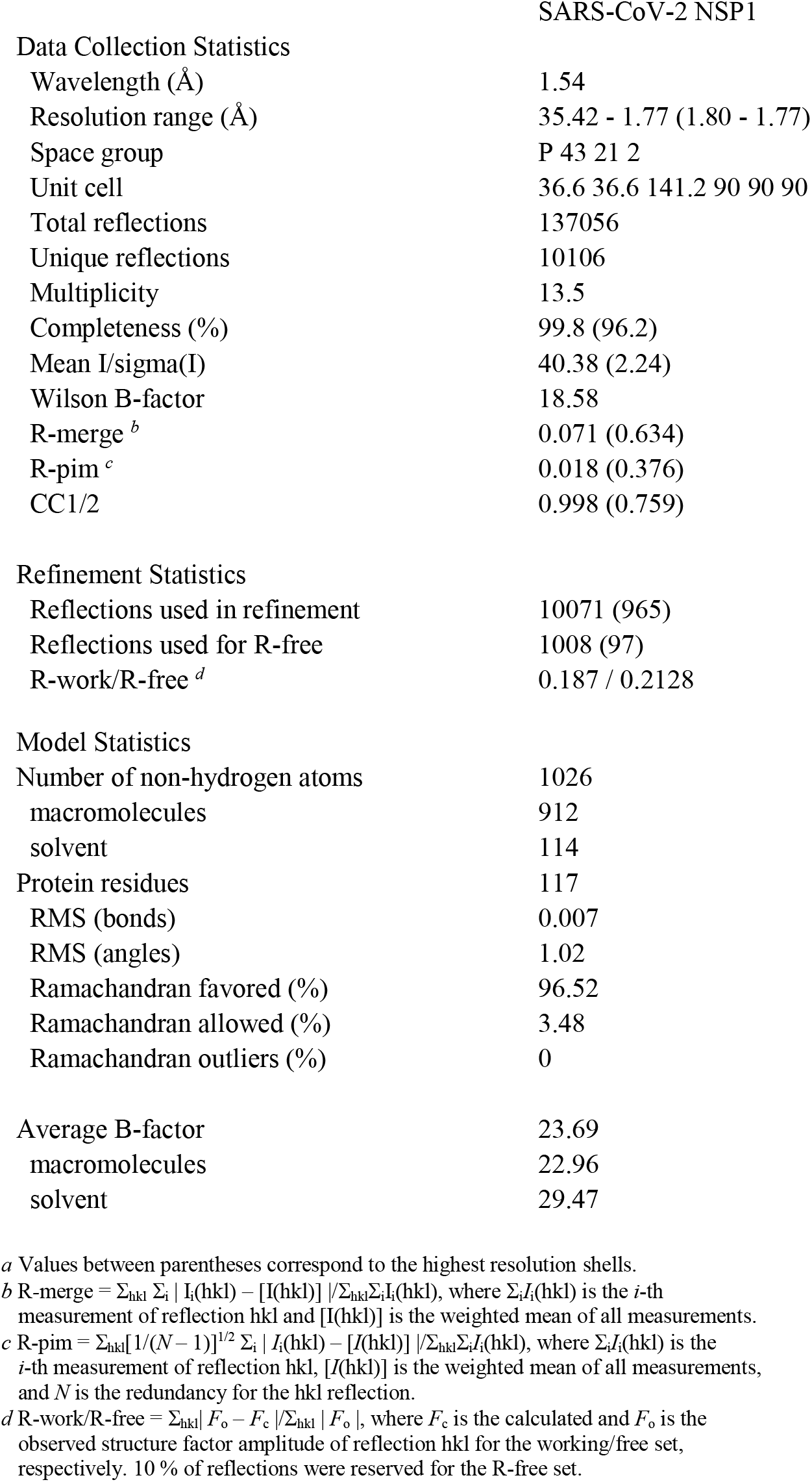
Data collection and Refinement Statistics.

### Comparison with solution structure of nsp1_13-127_ from SARS-CoV-1

The nsp1 portion of ORF1ab is well conserved when comparing SARS-CoV-1 and SARS-CoV-2 (84% identity, **(Fig. 2A**). This relatively high conservation is also applicable to the subset of amino acids (10-126) that comprise our structure (86%). This level of conservation between CoVs is remarkable given the limited amino acid sequence homology among nsp1s from different β-CoVs.(19) A number of mutations (9 of 16) occur in the large loops in the structure **(Fig. 2B)**. It is widely understood that loops often play critical roles in protein-protein interactions, making these mutations particularly interesting (24, 25).

**FIG 2.**
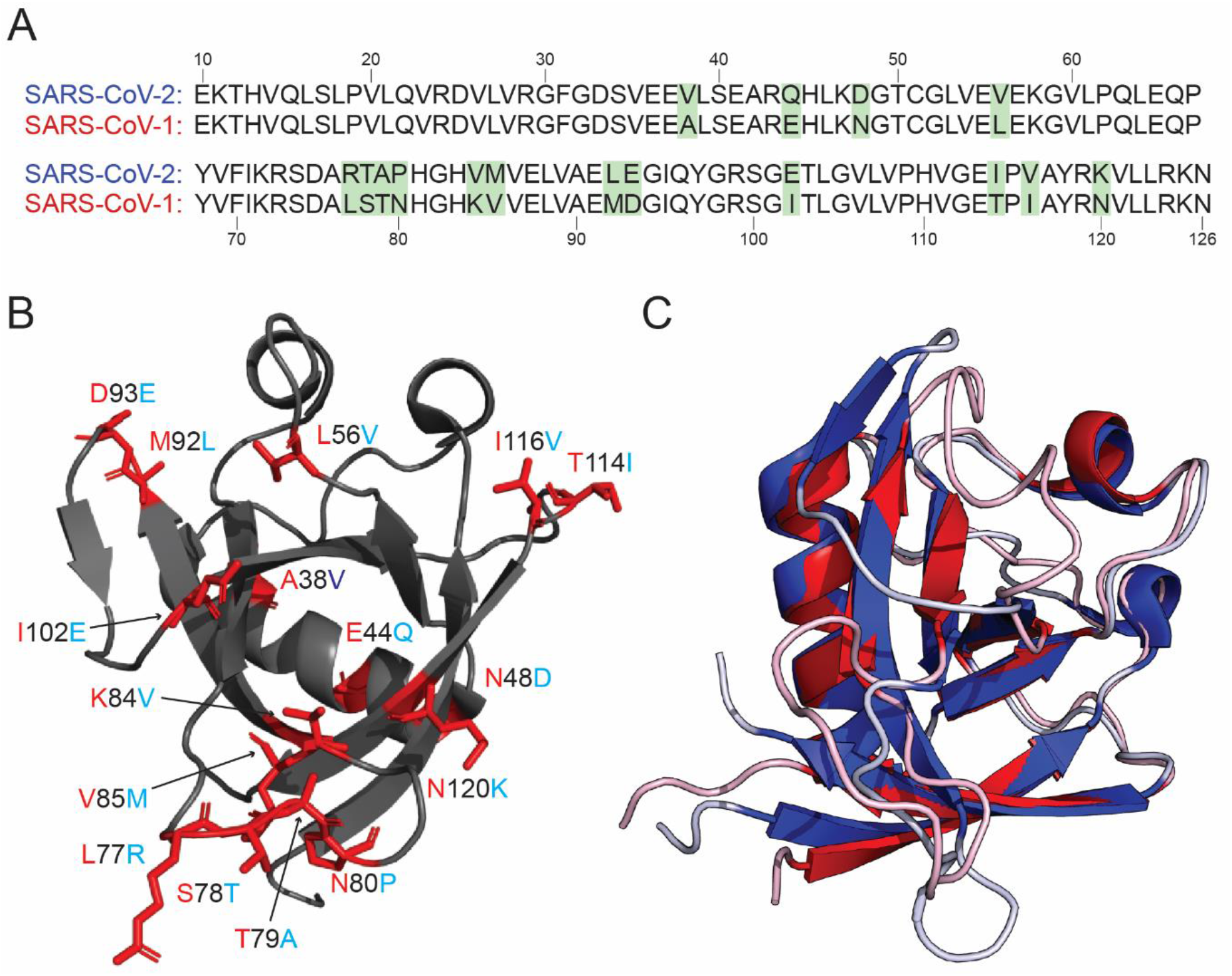
Comparison of the SARS-CoV-2 and SARS-CoV-1 nsp1 structures. (A) Sequence alignment between the nsp1 structures derived from SARS-CoV-2 and SARS-CoV-1 with amino acid differences highlighted in green. (B) Ribbon diagram of the SARS-CoV-2 nsp1_10-127_ with amino acid differences with SARS-CoV-1 nsp1_12-127_. The amino acid position is labeled in black with the sequence identity of SARS-CoV-1 at that position labeled in red and SARS-CoV-2 in light blue. (C) Overlay of the structures of nsp1 derived from SARS-CoV-1 (red) and SARS-CoV-2 (blue). PDB ID: 2HSX for SARS-CoV-1 nsp1_12-127_ and 7K7P for SARS-CoV-2 nsp1_10-127_.

We note that the structure is composed of approximately the same portions of nsp1 (residues 10-127) as the previously solved nsp1 structure from SARS-CoV-1 (residues 13-127). The fold of SARS-CoV-2 nsp1 is generally well conserved when compared to the previously solved structure of nsp1 from SARS-CoV-1. **(Fig. 2C)**. However, there are several notable differences between the structures upon detailed analysis. One such difference is the presence of an additional 3_10_ helix (η1) in the SARS-CoV-2 structure (residues 23-25). We note that the primary sequence composing this additional 3_10_ helix is identical between SARS-CoV-1 and SARS-CoV-2. Additionally, there are no apparent proximal mutations in SARS-CoV-2 that would stabilize this 3_10_ helix. The stabilization of this helix is due to a polar interaction between R24 in α1 and Q63 in α3 (**Fig. 3A**) present in only SARS-CoV-2 nsp1.

**FIG 3.**
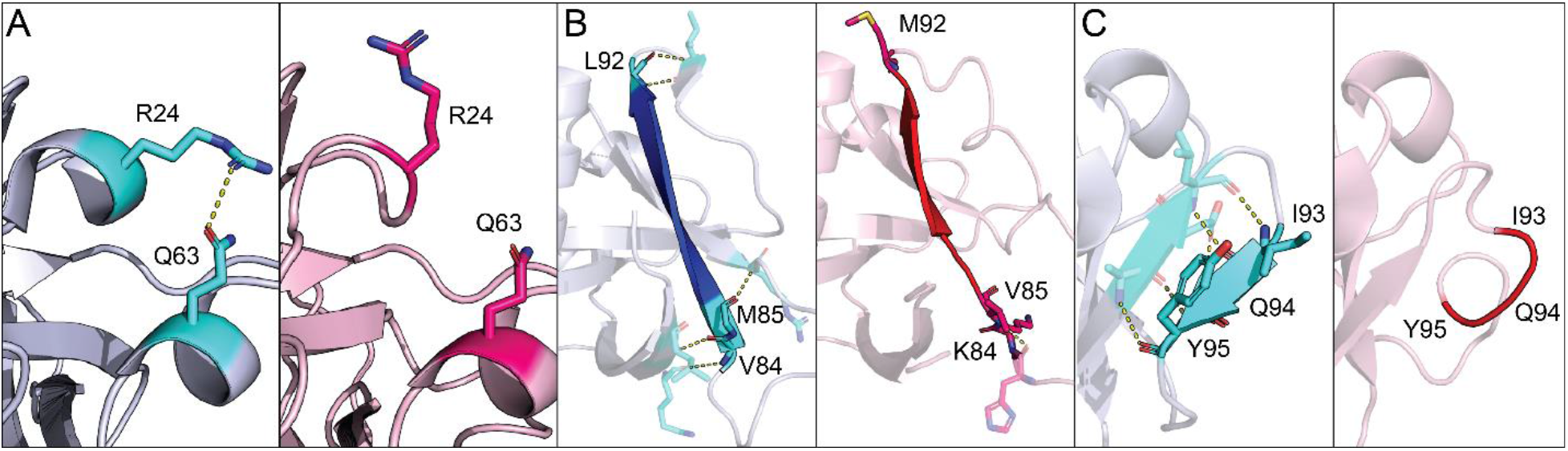
Differences in secondary structural elements between SARS-CoV-2 and SARS-CoV-1 nsp1. (A) Ribbon diagram of the stabilization via polar contacts of the 3_10_ helix present in SARS-CoV-2 nsp1 but not in SARS-CoV-1. Amino acids specified in the text are indicated in cyan and pink for SARS-CoV-2 and SARS-CoV-1, respectively. (B) Extension of the β4 strand in SARS-CoV-2 (blue) relative to SARS-CoV-1 (red). Mutations between the two viruses are labeled with their respective amino acid identity and position. Each mutation is also indicated in cyan for SARS-CoV-2 and pink for SARS-CoV-1. (C) An additional β-strand is present in the SARS-CoV-2 nsp1. Amino acids that compose this segment of both proteins are labeled and indicated in cyan for SARS-CoV-2 and pink for SARS-CoV-1. For all panels, SARS-CoV-2 nsp1 is in blue grey, SARS-CoV-1 nsp1 is in light pink, and polar contacts are indicated by a yellow dashed line.

In addition to the formation of a new 3_10_ helix in nsp1 from SARS-CoV2, there are also mutations that result in extension of β-strands within the interior of the protein when comparing the two proteins. The β4 strand is extended by four amino acids and is now composed by amino acids 84-92 in SARS-CoV-2 as opposed to amino acids 87-91 in SARS-CoV-1. This is due to the K84V, V85M, and M92L mutations in positions adjacent to the β4 strand **(Fig. 3B)**. This extension facilitates the formation of a new β-strand (β5) not found in the original SARS-CoV-1 structure. This strand is composed of amino acids 95-97 and is stabilized by main chain contacts with residues 90-92 **(Fig. 3C).** The β6 strand is also extended by two amino acids so it is now composed of amino acids 103-109 rather than amino acids 105-109. This extension is due to additional main chain interactions. Finally, β1, β3, and β6 (β5 in SARS-CoV-1) strands are also extended by a single amino acid.

One striking difference between the structures is the alternative conformations of two major loops located between β3 and β4 (loop 1) and between β4 and β6 (loop 2, β4 and β5 in SARS-CoV-1) **(Fig. 4).** The residues that make up these loops in SARS-CoV-2 nsp1 form a much more significant network of polar contacts with respect to those in the analogous loops in SARS-CoV-1 nsp1. These changes in local stability and conformation facilitate the formation of a new β-strand (β5) not found in SARS-CoV-1 nsp1. Analysis of loop 1 (residues 73-84) and loop 2 (residues 92-104) indicate an average Cα displacement of 16.6 Å and 27.6 Å, respectively. As loops play critical roles in protein-protein interactions, relative variations in these loops between the SARS-COV-1 and SARS-CoV-2 are particularly interesting as they may alter host-pathogen interactions involving nsp1 (24, 25).

**FIG 4.**
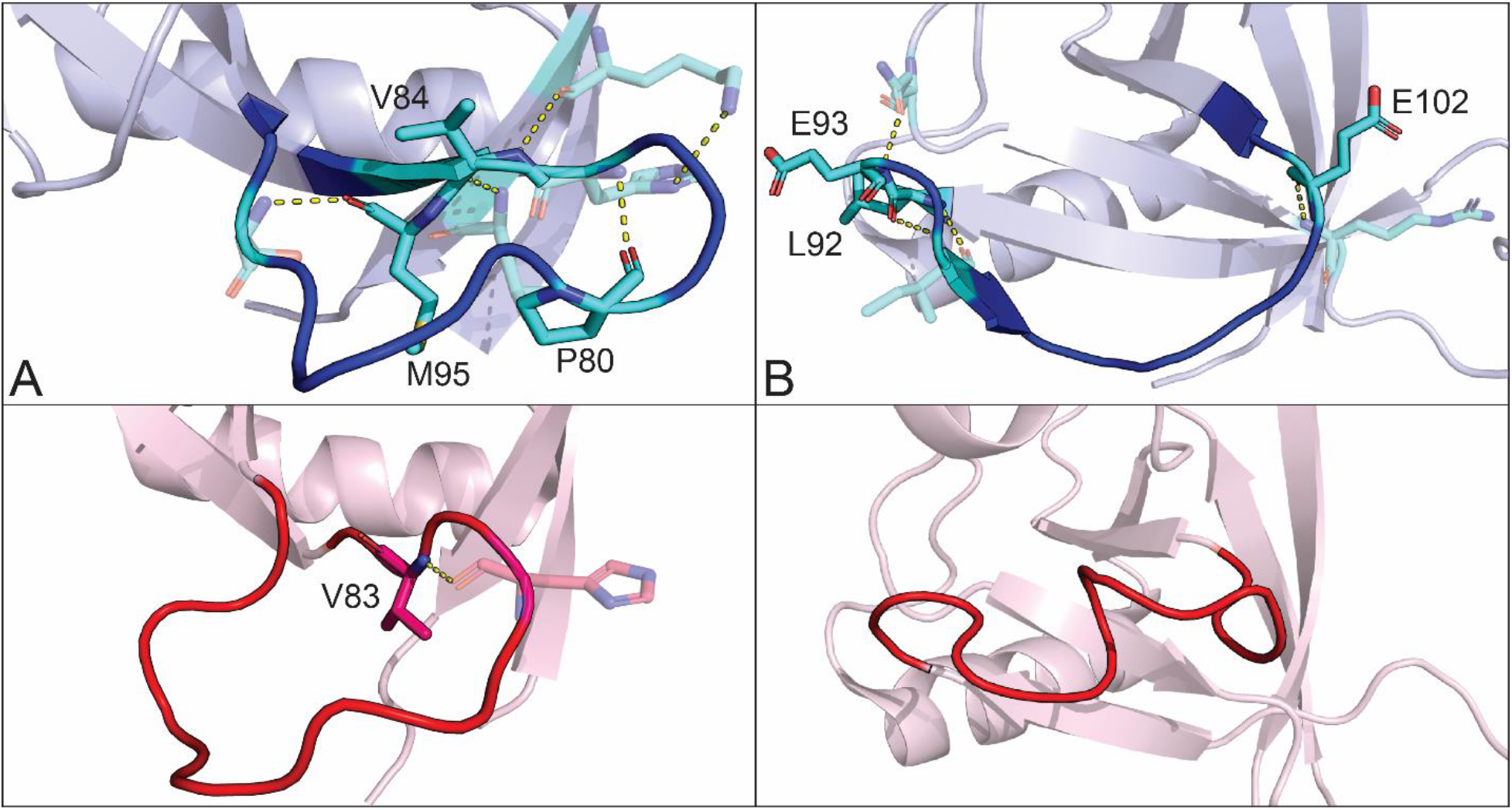
Conformational differences in major loops between SARS-CoV-2 and SARS-CoV-1 nsp1. Ribbon diagram of SARS-CoV-2 and SARS-CoV-1 nsp1 with the loops located between β3 and β4 (A) and β4 and β6 (B) are indicated in blue for SARS-CoV-2 and red for SARS-CoV-1. Residues mutated between the two viruses that are also involved in polar contacts are labeled as well as indicated in cyan for SARS-CoV-2 and pink for SARS-CoV-1. Polar contacts are indicated by a yellow dashed line.

Structural differences between the two proteins results in differences in the electrostatic surface potential. These differences are the result of conformational differences and mutations presented on the surface of the proteins. **Fig 5** contrasts four of these regions. Both SARS-CoV-1 and SARS-CoV-2 nsp1 share a large common patch of electronegative potential **(Fig. 5A and 5E)**, however in SARS-CoV-2 nsp1, the electronegative surface potential has an increased intensity and clustering has increased. This patch in SARS-CoV-2 nsp1 is generated by D33, E36, E37, E41, Q44, H45, D48, E65, Q66, and E93. These residues comprise three mutations from SARS-CoV-1 nsp1 and include E44Q, N48D, and D93E. In addition, a valley between the upper and lower portion of the patch in SARS-CoV-1 residues, between residues D33 and E65, is closed in SARS-CoV-2 by the reorientation of the main chain flanking Q66. This reorientation allows the sidechain of Q66 to form a new hydrogen bond with the amide nitrogen of E93. A dominant electropositive pocket and extended surface is also observed in both structures **(Fig. 5B and 5F)**. In SARS-CoV-2 nsp1, this region is composed of K11, R43, K47, R77, R124 and K125. The contiguous pit of electronegativity observed in SARS-CoV-2 nsp1 is disrupted in the SARS-CoV-1 structure by rearrangement of the C-terminus allowing G127 to protrude between R43 and K125. Two smaller juxtaposed electropositive and electronegative patches are shown in **Fig. 5C and 5G**. There are three mutations relative to SARS-CoV-1 nsp1 in this area: K84V, I102E, and N120K. In the SARS-CoV-2 structure, two mutations contribute to the negative surface character of this region. The mutated side chain of V84 occupies the space of K84 in SARS-CoV-1, while K120 buttresses the bottom of the patch. In the adjacent electropositive patch, E102 sits in the positive pocket surrounded by E55, E57, and E102. The conformation of the loop between β3 and β4, as well as the restructured strand (β5) and loop between β4 and β6 (β4 and β5 in SARS-CoV-1), contribute to vastly different surface contours **(Fig. 5C and 5G)**. Finally, a new electronegative crevasse is generated by the restructuring of strand β5 and the trailing loop leading to strand β6 and subsequent reposition of the concluding residues of β7 thru the C-terminus (residues 122 to 126) in SARS-CoV-2 **(Fig. 5D and 5H)**. Two prominent residues in this new crevasse are D75 and E87. These residues are identical in SARS-CoV-1, however the surrounding residues are positioned so that they are more obscured from the surface. E87 (SARS-CoV-1) is sandwiched between K72 and R124, appearing to salt bridge with K72.

**FIG 5.**
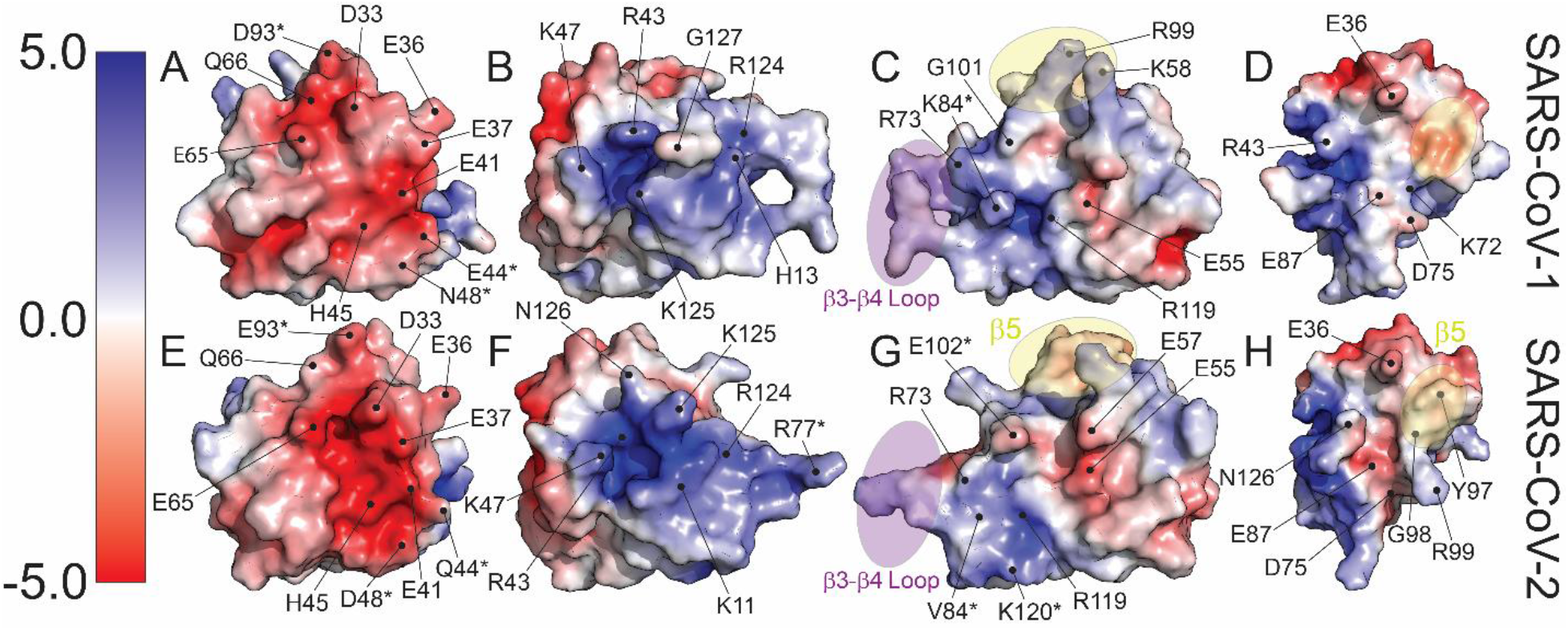
Differences in electrostatic surface potential between SARS-CoV-1 nsp1 and SARS-CoV-2 nsp1. Surface models of both proteins with amino acids contributing to the differences in electrostatic character labeled. Secondary structural elements that contribute to differences in surface contours are highlighted in purple for the loop in between β3 and β4 and in yellow for β5, found in SARS-CoV-2 but not SARS-CoV-1 nsp1. Residues that are mutated between the two viruses are indicated with an asterisk (*).

### Structural homology between SARS-CoV-2 nsp1 and α-CoV nsp1s

The structures of SARS-CoV-1 nsp1 and SARS-CoV-2 nsp1 represent the known nsp1 structures of β-CoVs. To date, the structures of five additional nsp1 structures originating from viruses in the α-coronavirus genera have been published. The sequence identity between SARS-CoV-2 nsp1 and these α-CoV is extremely low at 9-13%, with only marginally more sequence similarity at 21-27%. There is, however, easily identifiable structural homology between nsp1s derived from α- and β-CoVs. The structure of SARS-CoV-2 was structurally aligned with the structures of nsp1s from porcine transmissible gastroenteritis coronavirus strain Purdue (3ZBD)(26), porcine epidemic diarrhea virus (5XBC)(27), transmissible gastroenteritis virus (6IVC)(28), swine acute diarrhea syndrome coronavirus (6LPA)(29), and feline infectious peritonitis virus (6LP9)(29). Aligned structures are illustrated in **Fig. 6** with statistical analysis compiled in **Table 2**. All of the structures share a core topological similarity. However, there are substitutions and insertions in the linear arrangement of the secondary structure elements **(Fig. 6C)**. The first 3_10_ helix of SARS-CoV-2 structure is swapped for strand β2 in the α-CoV structures, though essentially occupying the same space. Strand β4 represents an insertion in the α-CoV structures. The strand forms a short parallel β-strand configuration with β2. As noted above, SARS-CoV-2 has new strand β5, not observed in SARS-CoV-1. This strand is also a deviation from the topology of the α-CoV nsp1 structures. The rest of the secondary structural elements follow a common fold and alignment being β1–α1–β2–η2–β3–β4–β6–β7 for SARS-CoV-2 and β1–α1–β3–η1–β5–β6–β7–β8 for α-CoVs **(Fig. 6H).**

**FIG 6.**
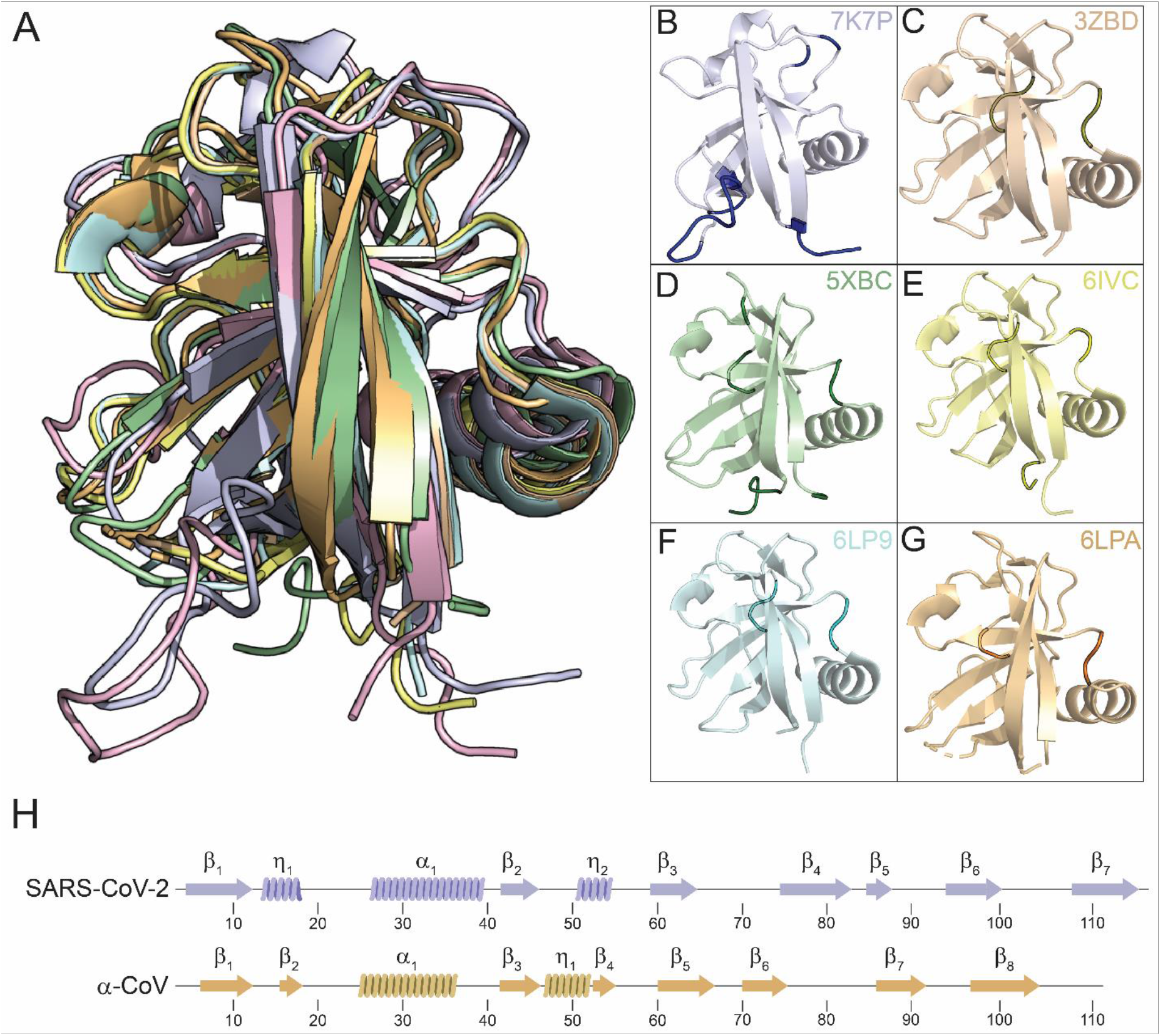
Structural Homology of β-and α-coronavirus nsp1s. (A) Structures of nsp1s from SARS-CoV-2 (7K7P), porcine transmissible gastroenteritis coronavirus strain Purdue (3ZBD), porcine epidemic diarrhea virus (5XBC), transmissible gastroenteritis virus (6IVC), swine acute diarrhea syndrome coronavirus (6LPA), and feline infectious peritonitis virus (6LP9) are shown structurally aligned in (A) and in common orientation (B)-(G). In (B)-(G), the common core shared with SARS-CoV-2 is shown in pale coloring, with deviations shown in darker shade coloring. (H) Secondary structural elements of SARS-CoV-2 and α-CoV nsp1 with positions indicated

**Table 2.**
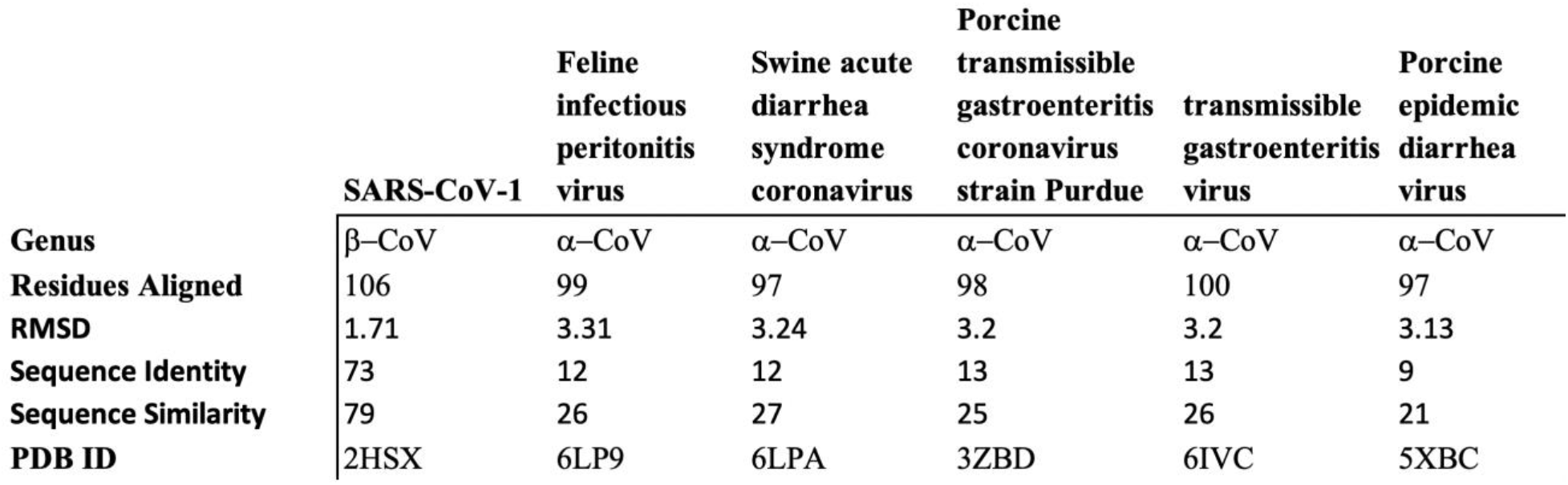
Structural Similarity of α-Coronavirus nsp1

## DISCUSSION

Here we report the high-resolution structure of nsp1 derived from SARS-CoV-2, the etiological agent of COVID-19. The interior of the protein is composed of a seven stranded β-barrel with an α-helix positioned on the side of the barrel and two 3_10_ helices across one end of the barrel. Nsp1 is highly homologous to SARS-CoV-1 nsp1 in both structure and sequence. However, there are a number of differences between the two proteins that may alter their function in the context of their respective viral life cycles. One difference is the presence of secondary structural elements, an additional β-strand (β5) and 3_10_ helix, in SARS-CoV-2 nsp1 that are not in SARS-CoV-1 nsp1. Several of the major loops in SARS-CoV-2 nsp1 are also in alternative conformations due to the increased polar contacts between amino acids that compose them and the globular domain. Finally, there are differences in the electrostatic surface potential resulting from amino acid differences between the two proteins. Given the critical role that nsp1 plays in the CoV lifecycle, these differences may underlie fundamental differences between the SARS-CoV-2 and SARS-CoV-2 such as virulence, pathogenicity, and infectivity.

Nsp1 is a virulence determinant of CoVs that plays critical roles during the viral lifecycle such as suppressing host gene expression (14, 30, 31). This suppression of host gene expression is necessary for viral replication and allows evasion of the cellular immune response. In a recent study, cryo-electron microscopy (cryo-EM) was used to show that the carboxyl terminus (residues 148-180) of SARS-CoV-2 nsp1 binds to 40S ribosomal subunit and inhibits translation (32, 33). Specifically, the structures revealed that this fragment of nsp1 binds to the mRNA entry channel of the 40S subunit which effectively blocks translation (32, 33). We modeled the structure of fulllength nsp1 using the iTasser platform with our structure as a template (34–36). As shown in Fig. 7, residues 1 – 9 and 131 – 162 contain no discernable secondary structure with the remaining residues adopting two α-helices. This is compatible with the cryo-EM structure that indicates that the carboxyl terminus (residues 148 – 180) binds to the 40S ribosomal subunit (32, 33).

**FIG 7.**
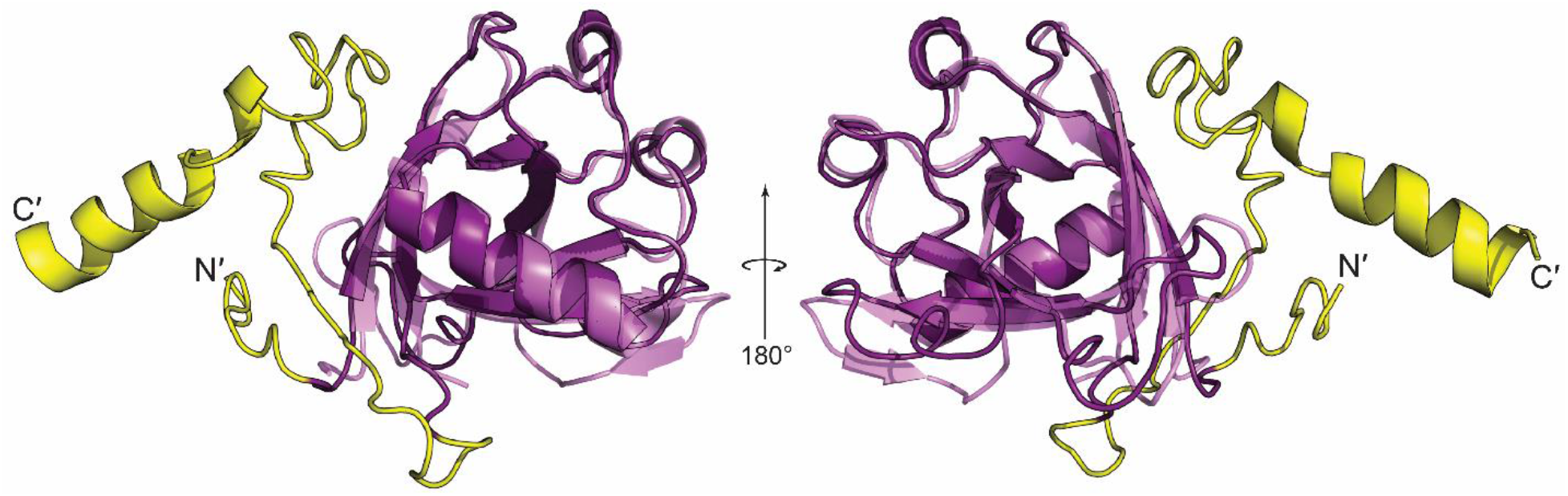
Structural prediction of full-length SARS-CoV-2 nsp1. Ribbon diagram of full-length SARS-CoV-2 nsp1 modeled using the iTasser platform.

Additionally, there is a high degree of structural and sequence homology between nsp1 derived from SARS-CoV-1 and SARS-CoV-2. The rare conservation amongst different β-CoVs may provide insight into SARS-CoV-2 nsp1 function from previous studies that focused on SARS-CoV-1 nsp1 (19). For example, previous studies have shown that nsp1 from SARS-CoV-1 is unique among CoVs and is hypothesized to contribute to the exceptional pathogenesis of SARS-CoV-1 in humans (37, 38). In fact, nsp1 control of interferon expression may be crucial to SARS-CoV-1 infection given that the virus is interferon-sensitive (15, 39, 40). Further studies will be needed to determine which functions are common between the two proteins and if there are any functional differences that alter their respective viral lifecycles.

There is an urgent need to develop both vaccines and antivirals against SARS-CoV-2 and any potential novel CoVs that pose a threat to global health. Because of its critical role, nsp1 has been identified as both a target for the development of broad-spectrum antivirals and rational live attenuated vaccine design (19, 20, 41–47). This fact was underscored by a previous study that focused on the function of nsp1 during the SARS-CoV-1 lifecycle which concluded the following: “How SARS-CoV nsp1 functions remains to be answered. Removal of the nsp1 protein function from the virus(48) will enable determination of its effects on replication and/or virulence *in vivo*; however, these experiments will require careful design and analysis because indiscriminate deletion of nsp1 sequence would remove *cis*-acting elements required for replication”(49). This conclusion is even more relevant during the COVID-19 pandemic. Our findings, combined with the structures of the carboxyl end of nsp1, are the first step to a comprehensive understanding of the structure-function relationship of SARS-CoV-2 nsp1.

## MATERIALS AND METHODS

### Protein expression and purification

The nsp1 segment of SARS-CoV-2 was subcloned into 6X His-SMT3 fusion T7 expression vector (pE-SUMO; Life Sensors) using primers to amplify residues 10-127 of ORF1a. Expression plasmids were transformed into DE3 Star BL21 *Escherichia coli* cells and grown in LB media containing 200 ug/mL of kanamycin. The nsp1 cultures were induced at OD_600_ of 0.7 with 1mM isopropyl β-D-1-thiogalactopyranoside for 48 h at 18 °C. Cells were lysed via sonication in buffer containing 20 mM HEPES, 500 mM NaCl, 5 mM Imidazole, 5 mM DTT and 1.3 mg/ml lysozyme at pH 7.8. Clarified lysates were loaded onto His-tag purification column (Roche) and the protein was eluted in a buffer containing 20 mM HEPES, 500 mM NaCl, 250 mM Imidazole, 2.5 mM DTT at pH 8. The 6X His-SMT3 tag was removed from the N terminus with recombinant SUMO protease. The cleaved sample was then reloaded onto the His-tag purification column to separate the 6X His-SMT3 tag from nsp1. Buffer exchange was performed with size exclusion chromatography using a Hi-Prep 26/60 Sephacryl S-100 HR (GE). The final buffer is composed of 20 mM NaPO4, 200 mM NaCl, 1 mM EDTA, 50 mM Arginine, 50 mM Glutamic Acid, 0.02% Sodium azide at a pH of 6.8.

### Crystallization, data collection, structure determination and refinement

Purified SARS-CoV-2 nsp1 was concentrated to 7 mg/mL and screened for crystallization conditions against commercially available kits using a Mosquito liquid handling robot (SPT Labtech). Diffraction quality crystals were obtained in conditions from multiple kits. An initial X-ray diffraction dataset was collected with the Pilatus 200K detector (Dectris) atop a MicroMax-007 HF rotating anode X-ray generator (Rigaku) from a crystal grown in 0.1 M MES monohydrate pH 6.5, 12% w/v polyethylene glycol 20,000, cryo-protected with a supplement of 20% glycerol. Raw intensity data were processed with the HKL3000 package.(50) Initial phases were solved by molecular replacement with PHASER(51) using the core structure of PDB ID: 2HSX. The structure was initially traced and refined with the AUTOBUILD routine in PHENIX(52), followed by manual editing in COOT(53) and refinement with phenix.refine. This nearly complete model was used as the model for molecular replacement with a final high-resolution dataset collected from a crystal grown in 0.2 M sodium acetate trihydrate, 0.1 M sodium cacodylate trihydrate pH 6.5, 30% w/v polyethylene glycol 8,000, cryo-protected with a supplement of 20% glycerol. The model was further traced manually and refined with phenix.refine. Data collection and refinement statistics are displayed in Table 1. All ribbon and surface illustrations of protein structures were prepared with PyMOL (Delano Scientific). Alignment statistics were determined with structure alignment tools from the RCSB (54).

## ACKNOWLEDGMENTS

This work was funded by the National Institutes of Health grants AI1346931 (C.M.P.) and AI116738 (T.J.G.) in addition to the UAB School of Medicine and Hugh Kaul Precision Medicine Institute COVID-19 Urgent Research Fund (C.M.P). The O’Neal Comprehensive Cancer’s X-ray crystallography core facility is supported through NIH P30CA13148.

